# High sensitivity cameras can lower spatial resolution in high-resolution optical microscopy

**DOI:** 10.1101/2024.02.23.581754

**Authors:** Henning Ortkrass, Marcel Müller, Anders Kokkvoll Engdahl, Gerhard Holst, Thomas Huser

## Abstract

High-resolution optical fluorescence microscopies and, in particular, super-resolution fluorescence microscopy, are rapidly adopting highly sensitive cameras as their referred photodetectors. Camera-based parallel detection facilitates high-speed live cell imaging with the highest spatial resolution. Here, we show that the drive to use ever more sensitive, photon-counting image sensors in cameras can, however, have detrimental effects on spatial resolution that many researchers are not aware of. Typical parameters which influence the selection of image sensors are pixel size, quantum efficiency, signal-to-noise performance, dynamic range, and frame rate of the sensor. A parameter that is, however, often overlooked, is the sensor’s modulation transfer function (MTF). We have determined the wavelength-specific MTF of front- and back-illuminated image sensors and evaluated how it affects the spatial resolution that can be achieved in high fluorescence microscopy modalities. We find significant differences in image sensor performance for sensors that cause the resulting spatial resolution to vary with up to 28%. This result shows that the choice of image sensor has significant impact on the imaging performance of all camera-based optical microscopy modalities.

## Introduction

The availability and application of image sensor based optical microscopy is rapidly increasing because of the high read-out rates combined with high quantum efficiency of modern scientific complementary metal-oxide semiconductor (sCMOS) based image sensors. The introduction of sCMOS sensors has also lowered the cost of ownership of single photon counting camera systems. High-speed readout of sCMOS cameras is paramount for advanced high-resolution live cell imaging, such as lattice light-sheet microscopy^1^, single-objective light sheet microscopy^2,3^ and oblique pane structured illumination microscopy^4^. sCMOS sensors employing rolling-shutter readout are now frequently being exploited in order to enable line-confocal out-of-focus signal rejection in novel light-sheet and super-resolution microscopy modalities, where the rolling shutter acts as a virtual pinhole slit^5,6^. These image sensors have further found rapid adoption in the majority of super-resolution imaging modalities, such as super-resolution structured illumination microscopy (SR-SIM)^7,8^, single molecule localization microscopy (SMLM)^9,10^, and signal fluctuation based super-resolution microscopies (e.g. SOFI)^11^, to name a few. Arguably, their most widespread use however is in standard wide-field fluorescence microscopy, where their performance is essential, especially in high-resolution applications ^12–14^.

sCMOS sensors can be realized in both front- and back-illuminated variants, and different semi-conductor parameters, e.g. resistance load. In front illumination, the electronic circuitry (transistors and conduits) are structured onto the light-sensitive front side of the sensor (the negative influence of the poor fill factor in many cases is partially compensated by the addition of micro lenses). In back-illuminated sensors, these structures are opposite of the light-sensitive surface. In general, this leads to a lower quantum efficiency (QE) of front-illuminated sensors, as by design some surface area is covered by the electronics and is not light sensitive. QE is often and rightfully seen as an important quality factor describing the sensitivity of an image sensor when choosing an image sensor for fluorescence microscopy applications, thus back-illuminated sensors typically appear as the superior choice for these systems.

Back-illumination of images sensors does, however, come with a drawback: once photons are converted to photo-electrons in the doped silicon, they have to traverse a much longer path through the thinned silicon to reach the potential well of the pixel that collects them. This increases the possibility of electrons being scattered into neighboring pixels, an effect called pixel crosstalk.^15^ While in principle known, this effect is often not accounted for, and most data-sheets quote QE as a key sensor parameter, but they do not provide the actual sensor’s modulation transfer function (MTF) that can be used to quantify this effect.

The incoherent wide-field detection scheme of the optical system of a fluorescence microscope itself, which is common to general wide-field imaging, such as wide-field fluorescence, SR-SIM, light-sheet microscopy, and spinning disk confocal microscopy, to name a few, has a MTF that falls off steeply towards higher spatial frequencies^16^. In a high-resolution optical system, where the magnification is chosen to achieve a projected pixel size that fulfills the Nyquist sampling criterion, the question arises whether or not a change in the camera sensor’s MTF actually contributes significantly to the overall resolution. And, if it does, will the higher QE provided by the back-illuminated sensors compensate for their lack in high-frequency response?

## Results and Discussion

### Modulation-transfer function (MTF) of different image sensors with the same pixel size

We investigated the effect of in total 4 different image sensors, two front-(C1 and C2)and two back-illuminated sCMOS sensors (C3 and C4), with the same pixel size of 6.5 μm x 6.5 μm and the same sensor size of 2048 x 2048 pixel (each in different camera implementations^*^) by determining their performance with wide field fluorescence and two popular super-resolution imaging methods, SR-SIM and direct stochastic optical reconstruction microscopy (dSTORM)^17^. A custom-constructed microscope, capable of high-resolution wide-field as well as SR-SIM imaging, was used for this purpose.The wide field images were acquired with a f=180 mm tube lens (overall magnification 60x) and a f=250 mm tube lens (overall magnification 83.3x) to measure the effect of the sensor MTF at two projected pixel sizes of 108 nm and 78 nm.^18^ Both fulfill the Nyquist criterion for a nominally 60x (f=3mm) 1.5NA objective lens at green emission. The super-resolution imaging was performed with a f=250 mm tube lens, thus 78 nm projected pixel size,and the fluorescence signal was split by a 50/50 beam splitter and focused on the two image sensors by identical tube lenses. Electronic synchronization allowed the two cameras to receive frame-by-frame identical data.

To quantify the effect of the MTF on super-resolution fluorescence images we first measured the overall MTF of the microscope for all image sensors. Single 100 nm TetraSpeck (TS) beads (Thermo Fisher Scientific) were imaged in the center of the field-of-view (FOV) ten times each, utilizing almost the maximum dynamic range of each camera. This was repeated with several different beads. The contrast of the images was maximized by subtracting the background and the image stacks were Fourier-transformed, deconvolved with the lateral projected spherical bead shape, and averaged^7^. The two-dimensional MTF was azimuthally averaged and set to zero outside its known support. The MTF was measured at 555 nm and 665 nm emission wavelength for front-illuminated (FSI) and back-illuminated (BSI) sensors. We confirmed that both, the transmitted and reflected image path after the beam splitter cube exhibited the same photon count rate and MTF (see Supplementary Information). In a well aligned microscope setup, it is assumed that the MTF mainly depends on the objective lens and the camera sensor - depending on the projected pixel size. We find that the MTF is different for all sensor types, especially for spatial frequencies higher than 1/μm at a projected pixel size of 78nm. The MTF of the BSI sensor C4 is, e.g. 24% lower than the MTF of the BSI sensor C3 at 665 nm and at a spatial frequency of 2/μm. This is explained by pixel crosstalk, which becomes more and more significant at higher spatial frequencies. The cutoff frequencies remain the same for all sensor types as these are only dependent on the numerical aperture of the objective lens. The actual image resolution, however, also depends on the magnitude of the MTF because Poisson noise limits the spectrum of the actually detectable spatial frequencies and, in general, contrast. To examine the effect of the different MTFs on wide field and super-resolved fluorescence images, we tested sensor C1 compared to C4 for wide field and SR-SIM, since they displayed the lowest and highest MTFs at 555 nm, and sensors C2 and C4 for dSTORM.

### Wide field imaging with 60x and 83.3x magnification

The camera sensors performance was first evaluated by imaging liver sinusoidal endothelial cells (LSEC) with different magnifications and the same optical setup (see Fig. 1). The magnification was changed by using different tube lenses with the same 60x 1.5NA Olympus objective lens. We used a standard 180 mm Olympus tube lens and a Ploessel type 250 mm tube lens to achieve fluorescence emission, the Rayleigh resolution limit is 216 nm, corresponding to a sampling of 2.0x at 60x magnification and a sampling of 2.77x at 83.3x magnification. Note that the 60x/180 mm objective lens/tube lens combination is used in a very large number of microscopes around the world. We measured the actual resolution achieved with the two sensors for these magnifications and at green emission by Fourier ring correlation on the wide field images. For the 60x magnification, five different FOVs were imaged ten times each, and the individual FRCs were averaged (Fig. 1c). With this modality, the images for the sensors were acquired sequentially at identical conditions. The front-illuminated sensor C1 gives a resolution limit of 242 nm and the back-illuminated sensor C4 310 nm. With the 83.3x magnification, two different FOVs were imaged 30 times on both sensors, simultaneously, and the averaged FRC (Fig. 1d) shows a resolution limit of 220 nm for C1 and 260 nm for C4. This result indicates that the image resolution is severely limited by the sensor MTF with a standard tube lens, and the resolution limit heavily depends on the sensor type. The effect is strong enough to not only be picked up by quantitative measurements such as FRC, but is also clearly visible by eye when comparing fine structural details (Fig. 1a,b magnified insets II vs IV).

**Figure 1:**
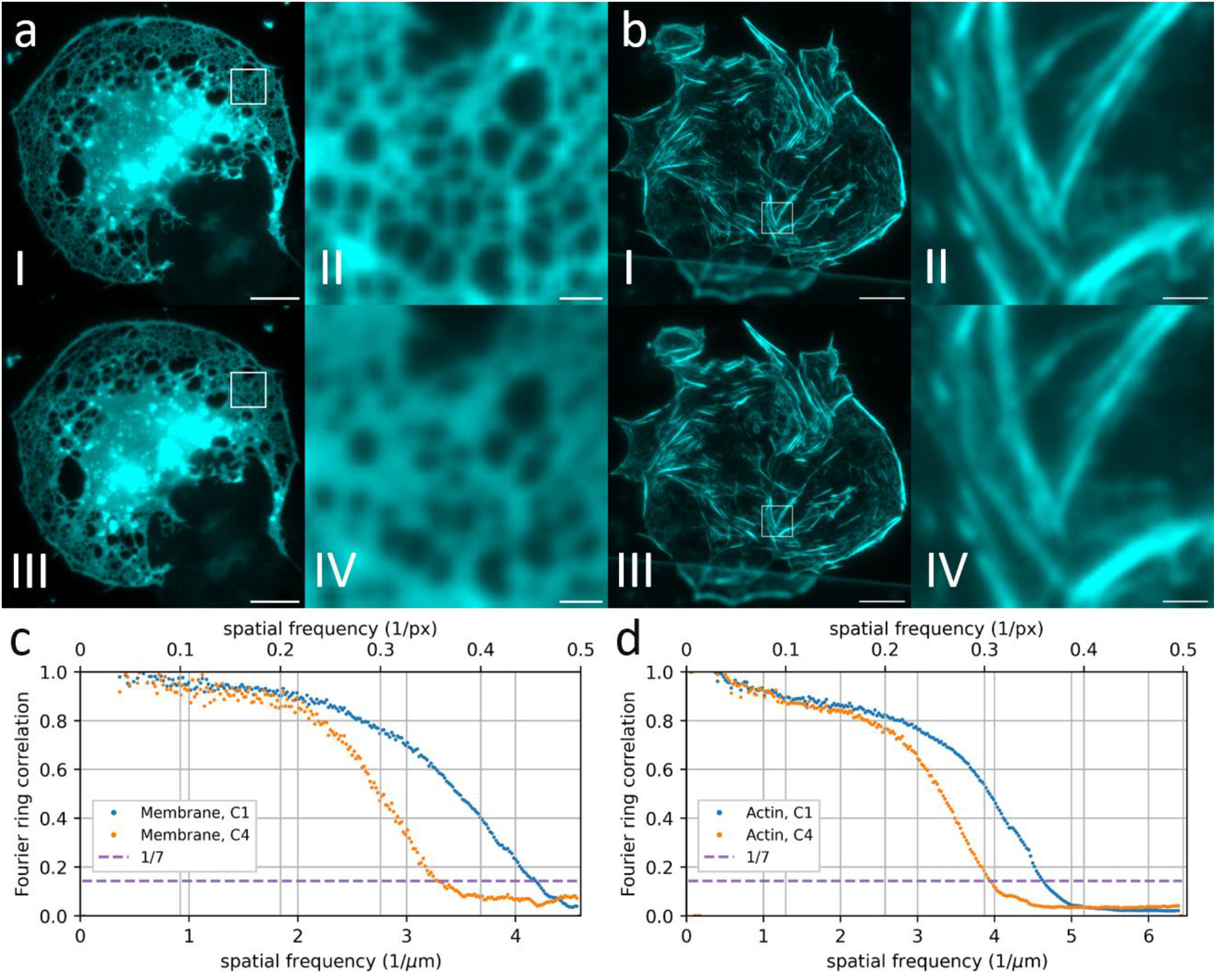
Wide-field images of liver sinusoidal endothelial cells stained with Vybrant DiO (a) and Phalloidin AF488 (b), both with green emission. The cells are imaged sequentially (a) or simultaneously (b) with the C1 sensor (aI, bI) and C4 sensor (aIII, bIII). To investigate the effect of the sensor MTF, different tube lenses with different magnifications were used. a was acquired with a total magnification of 60x, b with 83.3x. The images show a significant difference in resolution and contrast between the different sensors. The Fourier ring correlation (c) shows a sensor limited resolution limit of 242 nm for sensor C1 and 310 nm for C4 with 60x magnification.. It is calculated from the FRC average of six different FOV with 10 frames each. The FRC for the images with a 83.3x magnification (d) shows a resolution limit of 220nm for C1 and 260 nm for C4. It is the average of the FRC of two FOV with 30 frames each. Scale bar is 10 μm (aI, aIII, bI, bIII) and 1 μm (aII, aIV, bII, bIV).

With a magnification of 83.3x, the effect becomes less significant. This is to be expected, as the finer sampling (2.77x instead of 2.0x) spreads the PSF onto more pixels, so the effect of the sensor MTF becomes less pronounced in comparison. For super-resolution imaging, we thus chose the 83.3x magnification with the 250 mm tube lens, corresponding to a pixel size similar to commercial SR-SIM setups, where typically some oversampling is employed.

### Super-resolution structured illumination microscopy (SR-SIM)

The comparison of the camera sensors with respect to their performance in SR-SIM was tested on LSECs at 488 nm (stained against the actin cytoskeleton) and 640 nm (plasma membrane stain) in 2D- and total internal reflection fluorescence (TIRF) SIM mode, which result in different spatial resolutions^18^. Raw images were acquired simultaneously on both cameras and reconstructed with the same parameter set and the sensor-dependent MTF. The image reconstruction was performed using the open access fairSIM plugin in ImageJ^19,20^.

With a resolution improvement of ∼2x, which is typical for TIRF-SIM, we found a significant difference in the resolution limit for actin structures excited with 488nm and imaged at 505nm. The spatial resolution that we achieved with the C1 sensor (as measured by Fourier ring correlation (FRC)^21^) is 85nm, while the resolution limit of the images acquired with the C4 sensor is 93nm. which corresponds to a difference in resolution of 8% between the two different sensors. We explain this result based on the high frequency cutoff in the raw images of 4.9/μm which corresponds to 0.38/px and is close to the Nyquist sampling limit of 0.5/px and therefore most sensitive to pixel crosstalk. At lower frequency cutoffs, as obtained for the red-emitting plasma membrane stain, the FRC varies with less than 2%. Images acquired in 2D-SIM mode (resulting in a resolution improvement of ∼1.7x) show differences in the resolution limit between FSI and BSI sensors of less than 1% (Figure 2). This is explained by the lower spatial frequency of the excitation pattern, that causes a modulation pattern in the image data nearly unaffected by pixel crosstalk.

**Figure 2:**
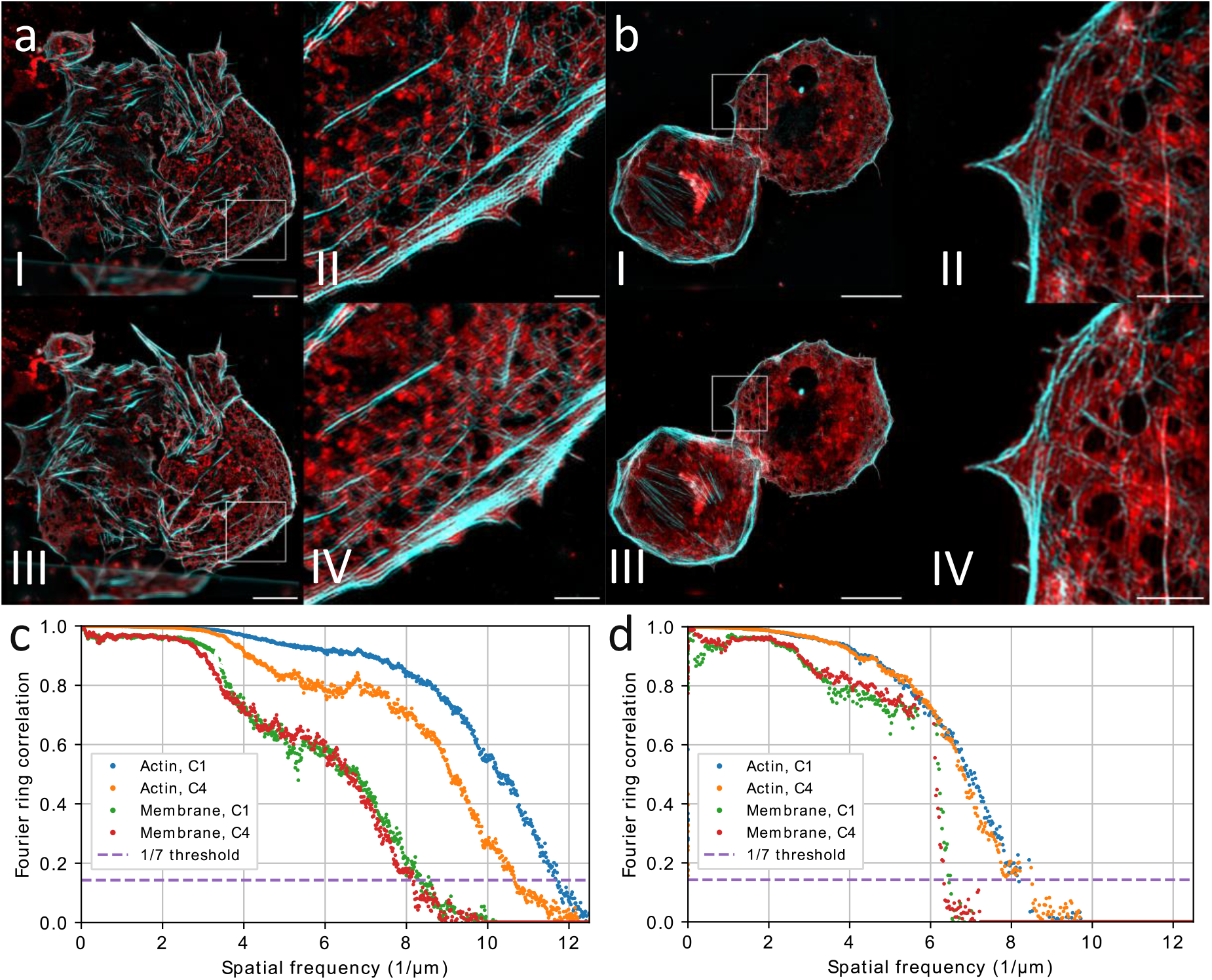
Reconstructed SR-SIM images of liver sinusoidal endothelial cells, stained with Phalloidin AF488 and membrane dye Biotracker 647. The cells are imaged simultaneously by the C1 sensor (aI, bI) and the C4 sensor (aIII, bIII) in TIRF (a) and 2D-SIM (b) mode. Fourier ring correlation (FRC) analysis reveals the different resolution improvements for TIRF-SIM (c) and 2D-SIM (d) for both sensor types and both wavelengths. The resolution limit corresponding to the frequency cutoff is significantly different for the actin cytoskeleton (show in cyan) imaged by excitation at 488 nm with TIRF-SIM: 85nm with the C1 sensor and 93nm with the C4 sensor. The FRC curves vary also for the plasma membrane (excited at 647 nm) imaged in TIRF and 2D-SIM mode, but at red wavelengths the resolution limit varies less than 2% for the sensor types. Scale bar is 10 μm (aI, aIII, bI, bIII) and 2 μm (aII, aIV, bII, bIV).

### Direct stochastic optical reconstruction microscopy (dSTORM)

In order to investigate the influence of MTF’s of the different sensor types on single molecule localization microscopy (SMLM), we performed dSTORM on immunofluorescently labeled microtubules in U2OS cells. The same microscope setup, in particular the same detection scheme, was used as in the SR-SIM experiments. However, instead of SIM patterns, classic widefield fluorescence excitation at 647 nm wavelength, was used with the fluorescence signal again split 50:50 to the two sCMOS image sensors. As can be seen in Fig. 3, the dSTORM reconstruction of 10,000 camera frames exhibits only a slight difference in spatial resolution between data acquired by the different image sensors. This is expected based on the wavelength-specific response of the sensor and further exacerbated by the dSTORM image reconstruction process, where the image is composed of points representing the centroid of a 2D fit function for each molecule detected. The precision of localization scales with the PSF width, which is somewhat influenced by the sensor MTF, but also with the square root of the number of detected photons, which of course benefits from an increase in sensor quantum efficiency. The effect of the MTF on dSTORM localization precision therefore appears to be negligible for this type of super-resolution microscopy.

**Figure 3:**
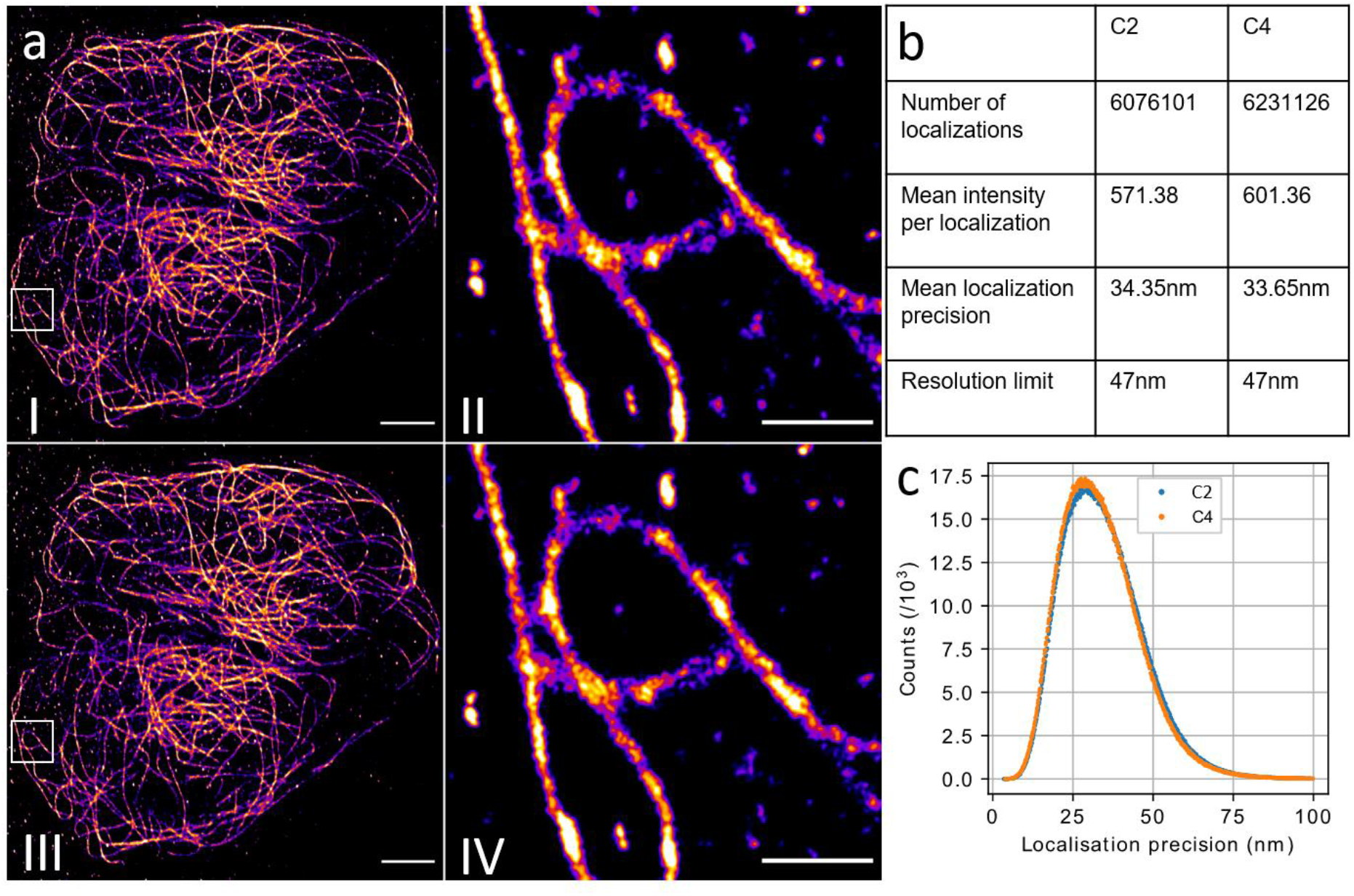
Super-resolution direct stochastic optical reconstruction microscopy images of Alexa647-immunofluorescently stained microtubuli in an U2OS cell (a). (aI, aII): reconstruction of the raw data acquired by the C2 sensor, (aIII, aIV): simultaneously acquired images using the C4 sensor, reconstructed with the same parameters. The mean localization precision (c) and resolution is similar for both cameras, despite the higher photon count per localization obtained with the back-illuminated sensor. Scale bars are 5μm (aI), (aIII) and 1μm (aII), (aIV).

## Conclusions

We have demonstrated that the image sensor type has a significant impact on a microscope’s MTF and therefore significantly affects the spatial resolution in wide field fluorescence imaging close to the Nyquist limit.

At 60x magnification a very popular choice for many commercially available implementations of high-NA, high-resolution wide-field and super-resolution imaging, the effect significantly reduces the image quality. A similar impact should be expected for any other high-resolution imaging task where pixel size is kept close to or even below the Nyquist criterion.

At 83.3x magnification, a choice providing some Nyquist oversampling (typical for commercial high- and super-resolution systems), the effect is less pronounced but still clearly noticeable in both wide-field and SR-SIM imaging modalities. For dSTORM, as an example of SMLM, however, it is not significant, as the reliance on molecule localization and their dependence on detected photons compensates for the reduced MTF.

The MTFs of the image sensors differ significantly for spatial frequencies in the image plane above 0.1/pixel. We explain this effect with the underlying sensor architecture, which affects the likelihood of pixel crosstalk. This affects both wide-field imaging as well as TIRF-SIM reconstructions at emission wavelengths in the green part of the visible spectrum with a frequency cutoff close to the Nyquist sampling rate. The MTF directly impacts the signal-to-noise ratio of high spatial frequencies of the reconstructed image and thus the resolution limit. Here, sample structures imaged at green wavelengths exhibit a difference in spatial resolution of up to 28%. In summary, the MTF of the image sensor plays a critical role in the ultimate spatial resolution that can be achieved in high resolution microscopy with modern sCMOS image sensors. This result also impacts other types of camera-based high resolution microscopy modalities, such as spinning-disk confocal,light-sheet fluorescence, and even electron microscopy. Depending on the application it should be carefully evaluated which sensor type is chosen, because the sensor type (FSI vs. BSI) can play a more critical role than the quantum efficiency of the sensor.

## Acknowledgements

The authors would like to thank Prof. Peter McCourt and Dr. Karolina Szafranska (UiT - the Arctic University of Norway) for the kind gift of rat liver sinusoidal endothelial cells. This work received funding from the European Union’s European Innovation Council PATHFINDER Open Programme under grant agreement No 101046928.

## Footnotes

* C1: pco.edge 4.2 (FSI image sensor: CIS2020AF), C2: pco.panda 4.2 (FSI image sensor: GSENSE2020), C3: pco.edge 4.2 bi (BSI image sensor: GSENSENE2020BSI-H), C4: pco.panda 4.2 bi (BSI image sensor: GSENSE2020BSI-M)

## References

1. Chen, B.-C. et al. Lattice light-sheet microscopy: Imaging molecules to embryos at high spatiotemporal resolution. Science 346, 1257998 (2014).

2. Bouchard, M. B. et al. Swept confocally-aligned planar excitation (SCAPE) microscopy for high-speed volumetric imaging of behaving organisms. Nat. Photonics 9, 113–119 (2015).

3. Voleti, V. et al. Real-time volumetric microscopy of in vivo dynamics and large-scale samples with SCAPE 2.0. Nat. Methods 16, 1054–1062 (2019).

4. Chen, B. et al. Resolution doubling in light-sheet microscopy via oblique plane structured illumination. Nat. Methods 19, 1419–1426 (2022).

5. Baumgart, E. & Kubitscheck, U. Scanned light sheet microscopy with confocal slit detection. Opt. Express 20, 21805–21814 (2012).

6. Schropp, M., Seebacher, C. & Uhl, R. XL-SIM: Extending Superresolution into Deeper Layers. Photonics 4, 33 (2017).

7. Gustafsson, M. G. L. et al. Three-Dimensional Resolution Doubling in Wide-Field Fluorescence Microscopy by Structured Illumination. Biophys. J. 94, 4957–4970 (2008).

8. Schermelleh, L. et al. Subdiffraction Multicolor Imaging of the Nuclear Periphery with 3D Structured Illumination Microscopy. Science 320, 1332–1336 (2008).

9. Betzig, E. et al. Imaging intracellular fluorescent proteins at nanometer resolution. Science 313, 1642–1645 (2006).

10. Huang, B., Wang, W., Bates, M. & Zhuang, X. Three-dimensional Super-resolution Imaging by Stochastic Optical Reconstruction Microscopy. Science 319, 810–813 (2008).

11. Dertinger, T., Colyer, R., Iyer, G., Weiss, S. & Enderlein, J. Fast, background-free, 3D super-resolution optical fluctuation imaging (SOFI). Proc. Natl. Acad. Sci. 106, 22287–22292 (2009).

12. Oliinyk, O. S. et al. Deep-tissue SWIR imaging using rationally designed small red-shifted near-infrared fluorescent protein. Nat. Methods 20, 70–74 (2023).

13. Skrabak, D. et al. Slack K+ channels limit kainic acid-induced seizure severity in mice by modulating neuronal excitability and firing. Commun. Biol. 6, 1–12 (2023).

14. Gross, D. et al. IKCa channels control breast cancer metabolism including AMPK-driven autophagy. Cell Death Dis. 13, 1–14 (2022).

15. Estribeau, M. & Magnan, P. CMOS pixels crosstalk mapping and its influence on measurements accuracy in space applications. In Sensors, Systems, and Next-Generation Satellites IX vol. 5978 315–326 (SPIE, 2005).

16. Manton, J. D., Ströhl, F., Fiolka, R., Kaminski, C. F. & Rees, E. J. Concepts for structured illumination microscopy with extended axial resolution through mirrored illumination. Biomed. Opt. Express 11, 2098–2108 (2020).

17. Heilemann, M., van de Linde, S., Mukherjee, A. & Sauer, M. Super-resolution imaging with small organic fluorophores. Angew. Chem. Int. Ed Engl. 48, 6903–6908 (2009).

18. Ortkrass, H. et al. High-speed TIRF and 2D super-resolution structured illumination microscopy with large field of view based on fiber optic components. Opt. Express in print, (2023).

19. Müller, M., Mönkemöller, V., Hennig, S., Hübner, W. & Huser, T. Open-source image reconstruction of super-resolution structured illumination microscopy data in ImageJ. Nat. Commun. 7, 10980 (2016).

20. Schneider, C. A., Rasband, W. S. & Eliceiri, K. W. NIH Image to ImageJ: 25 years of image analysis. Nat. Methods 9, 671–675 (2012).

21. Nieuwenhuizen, R. P. J. et al. Measuring image resolution in optical nanoscopy. Nat. Methods 10, 557–562 (2013).

