## Supplementary file for "High sensitivity cameras can lower spatial resolution in high-resolution optical microscopy"

Henning Ortkraß[1], Marcel Müller[1], Anders Kokkvoll Engdahl[1], Gerhard Holst[2], and Thomas Huser[1]
[1]Biomolecular Photonics, Faculty of Physics, Bielefeld University, 33615 Bielefeld, Germany
[2]Excelitas PCO GmbH, Donaupark 11, 93309 Kelheim, Germany

### Setup

The MTF measurements and SIM-imaging experiments were performed with a wide-field microscope with a fiber-based 2D-SIM excitation path. We used a 491nm laser for excitation (Cobolt Calypso 100), as well as a 532nm laser (Cohorent Compass 215M-50) and a 639nm laser (Photontec MSL-FN-639-300). The excitation pattern was projected into the sample with an Olympus 60x 1.5NA objective lens (UPLAPO60XOHR) and the fluorescence was epi-detected, split by a 50/50 non-polarizing beam splitter cube (Qioptiq G335525000) and imaged by a Ploessel-type tube lens (constructed out of two Thorlabs achromatic lenses AC508-500-A) onto the image sensors. The difference in the MTF between both imaging paths was checked to be negligible. The projected pixel size was 78nm.
The camera abbreviations correspond to the following camera and sensor models:

| | Camera model | Sensor type |
|---|---|---|
| C1 | pco.edge 4.2 | FSI image sensor: CIS2020AF |
| C2 | pco.panda 4.2 | FSI image sensor: GSENSE2020 |
| C3 | pco.edge 4.2 bi | BSI image sensor: GSENSENE2020BSI-H |
| C4 | pco.panda 4.2 bi | BSI image sensor: GSENSE2020BSI-M |

### MTF measurement

For the calculation of the systems MTF for different sensors, we imaged single 100nm TetraSpeck beads (ThermoFisher T7279) within a field of view of $256x256px^2$ ten times. The stack was Fourier-transformed, reduced to real-values and averaged. The resulting image was azimuthally averaged. This protocol was performed twice for different beads and the resulting MTFs were averaged. The final MTF was deconvolved with the 2D-projected shape of the spherical bead.

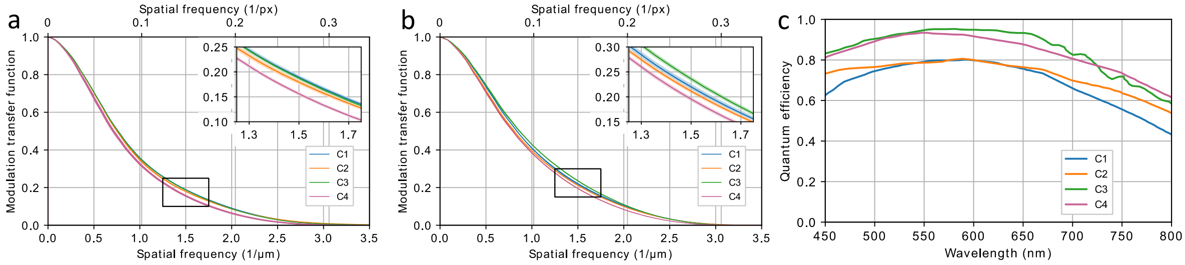


***Figure 1**:* Comparison of the modulation transfer function (MTF) of the front-illuminated (FSI) and back-illuminated (BSI) image sensors**.** The sensor dependent modulation transfer function (MTF) of the setup is measured at 555nm (a) and 665nm (b) emission wavelength. The spatial frequency in 1/px corresponds to the camera plane, the spatial frequency in 1/µm to the sample plane. The difference of the MTF of the BSI sensor compared to the FSI sensors is most significant between 1.5/µm and 2/µm. The quantum efficiency (c), that also affects the quality of super-resolved image reconstruction, is higher for the BSI sensor types.

### Cell preparation

U2OS cells are seeded on #1.5 glass cover slips. For indirect antibody-staining of tubulin filaments, the U2OS cells are fixed with 4% paraformaldehyde for 10 minutes at room temperature. An extraction step is followed by fixation with 0.5% glutaraldehyde in PEM buffer (PEM: 80mM piperazine-N,N-bis(2-ethanesulfonic acid) (PIPES), 5mM egtazic acid (EGTA), 2mM $MgCl_2$ at pH 6.8). After washing with phophate buffered saline (PBS), glutaraldehyde-induced autofluorescence is quenched by addition of 0.1% $NaBH_4$ in PBS for 7 minutes followed by washing with PBS three times.

Cells are blocked and permeabilized using a blocking buffer containing 0.3% gelatin and 0.05% Triton X-100 in PBS for 1 hour. Staining is performed overnight at 4 °C for $\alpha$- and $\beta$-tubulin using a mixture of three primary antibodies (T5168, T6199, T5923, Sigma) at a combined dilution of 1:150 in blocking buffer. Cells are washed with PBS three times and the second staining solution, AF647-conjugated goat anti-mouse IgG secondary antibody (A-21237, ThermoFisher) diluted 1:200 in blocking buffer, is incubated at room temperature for 90 minutes. The common GODCAT buffer containing the enzymatic oxygen scavengers glucose oxidase and catalase, with beta-mercaptoethanol (BME) as switching agent was used to induced intermittent fluorescence of the AF647 fluorophores.

Cryo-preserved rat liver sinusoidal endothelial cells (LSECs) are a kind gift from Dr. Peter McCourt and Dr. Karolina Szafranska at UiT - the Arctic university of Norway. They were shipped on dry ice to Germany and stored at -80°C. For thawing and seeding, a vial with LSECs is placed in an incubator at 37°C until nearly all the ice is thawed. The cells are gently pipetted drop-wise to 25 ml of pre-warmed Dulbecco's Modified Eagle Medium (DMEM) and centrifuged at 50g for 3 minutes to remove any hepatocytes remaining from the cell isolation. The supernatant containing LSECs is used for a second centrifugation step at 300g for 8 minutes. The cell pellet is resuspended in 4 ml - 7 ml DMEM and 1.5 ml (~100.000 cells per $cm^2$) of the cell solution is pipetted onto a fibronectin-coated #1.5 glass coverslip. The coverslip surface is coated with fibronectin (0.2 mg/ml) in phosphate buffered saline (PBS) containing 2 mM ethylenediaminetetraacetic acid (EDTA) for 1 hour at room temperature and washed with PBS afterwards. After allowing the cell suspension to incubate on the glass coverslip for 1 hour at 37°C and 5% $CO_2$, the coverslip was washed with pre-warmed DMEM and incubated for another 2 hours before fixation with 4% formaldehyde in PBS for 10 minutes at room temperature. First the plasma cell membrane is stained with BioTracker 655 Red Cytoplasmic Membrane Dye (SCT108, Sigma) diluted 1:200 in PBS for 1 hour at room temperature. The cells were washed twice in PBS before staining the actin cytoskeleton. The LSECs were incubated in a 1:40 dilution of Phalloidin CF568 (00044-T, Biotium) in PBS for 2 hours at room temperature. After the staining process was completed, the cells were washed 3 times with PBS.
